# Specificity and Mechanism of Coronavirus, Rotavirus and Mammalian Two-Histidine-Phosphoesterases That Antagonize Antiviral Innate Immunity

**DOI:** 10.1101/2021.06.16.448777

**Authors:** Abhishek Asthana, Christina Gaughan, Susan R. Weiss, Robert H. Silverman

**Affiliations:** Department Cancer Biology, Cleveland Clinic Foundation, Lerner Research Institute, Cleveland, OH, 44195, United States; Department of Microbiology, Perlman School of Medicine at the University of Pennsylvania, Philadelphia, PA, 19104, United States; Penn Center for Research on Coronaviruses and Other Emerging Pathogens, Perlman School of Medicine at the University of Pennsylvania, Philadelphia, PA, 19104, United States

## Abstract

2’,5’-oligoadenylate(2-5A)-dependent endoribonuclease, RNase L, is a principal mediator of the interferon (IFN) antiviral response. Therefore, regulation of cellular levels of 2-5A is a key point of control in antiviral innate immunity. Cellular 2-5A levels are determined by IFN-inducible 2’,5’-oligoadenylate synthetases (OASs) and by enzymes that degrade 2-5A. Importantly, many coronaviruses and rotaviruses encode 2-5A degrading enzymes thereby antagonizing RNase L and its antiviral effects. A-kinase anchoring protein 7 (AKAP7), a mammalian counterpart, could possibly limit tissue damage from excessive or prolonged RNase L activation during viral infections or from self double-stranded-RNAs that activate OAS. We show these enzymes, members of the two-histidine-phosphoesterase (2H-PE) superfamily, constitute a sub-family referred here as 2’,5’-PEs. 2’,5’-PEs from mouse coronavirus (CoV) MHV (NS2), MERS-CoV (NS4b), group A rotavirus (VP3), and mouse (AKAP7) were investigated for their evolutionary relationships and activities. While there was no activity against 3’,5’-oligoribonucleotides, all cleaved 2’,5’-oligoadenylates efficiently, but with variable activity against other 2’,5’-oligonucleotides. The 2’,5’-PEs are shown to be metal ion-independent enzymes that cleave trimer 2-5A (2’,5’-p_3_A_3_) producing mono- or di- adenylates with 2’,3’-cyclic phosphate termini. Our results suggest that elimination of 2-5A might be the sole function of viral 2’,5’-PEs, thereby promoting viral escape from innate immunity by preventing or limiting the activation of RNase L.

**IMPORTANCE:** Viruses often encode accessory proteins that antagonize the host antiviral immune response. Here we probed the evolutionary relationships and biochemical activities of two-histidine-phosphoesterases (2H-PEs) that allow some coronaviruses and rotaviruses to counteract antiviral innate immunity. In addition, we investigated the mammalian enzyme, AKAP7, which has homology and shared activities with the viral enzymes and might reduce self-injury. These viral and host enzymes, that we refer to as 2’,5’-PEs, specifically degrade 2’,5’-oligoadenylate activators of the antiviral enzyme RNase L. We show that the host and viral enzymes are metal ion independent and exclusively cleave 2’,5’- and not 3’,5’-phosphodiester bonds, producing cleavage products with cyclic 2’,3’-phosphate termini. Our study defines 2’,5’-PEs as enzymes that share characteristic conserved features with the 2H-PE superfamily but which have specific and distinct biochemical cleavage activities. These findings may eventually lead to pharmacologic strategies for developing antiviral drugs against coronaviruses, rotaviruses, and other viruses.

## INTRODUCTION

How interferons (IFNs) inhibit viral infections, and how viruses antagonize the IFN antiviral response, have been investigated for the past few decades, but with renewed intensity as a result of the SARS-CoV-2 pandemic (1–4). Mammalian cells often detect and respond to viruses after sensing viral double-stranded (ds)RNA, a common viral pathogen associated molecular pattern (PAMP) that induces type I and type III IFNs (1, 2). These IFNs induce expression of hundreds of IFN stimulated genes (ISGs), including numerous antiviral effector proteins (5). Included among the human antiviral proteins encoded by ISGs are 2’,5’-oligoadenylate (2-5A) synthetases 1-3 (OAS1-3) consisting of one, two and three core OAS units, respectively (6–8). However, not all mammalian species express a similar set of homologous OASs, and some, but not all, related OASL proteins lack enzymatic activity (9, 10). Upon binding of, and activation by, viral dsRNA OAS1-3 synthesize 2-5A [p_3_(A2’p5’)_n_A, where n=2 to >3] from ATP (7). The only known function of 2-5A is dimerization and activation of RNase L resulting in degradation of host and viral RNA, cessation of protein synthesis, apoptosis and inflammasome activation (11–15)(Fig. 1A). In addition to its antiviral effects, RNase L resulted in cell death in response to mutation of ADAR1 (adenosine deaminase acting on RNA-1) in a cell line or in cells treated with the DNA demethylating drug 5-aza-cytidine, both of which induce synthesis of self-dsRNA from repetitive DNA elements in the genome (16–18). Thus, regulation of 2-5A levels is critical for host cell viability as well as for control of viral infections and pathogenesis. Yet there are gaps in our knowledge of precisely how levels of 2-5A are established to restrict viral replication and spread by RNase L activation while at the same time minimizing tissue damage to the host.

**Figure 1.**
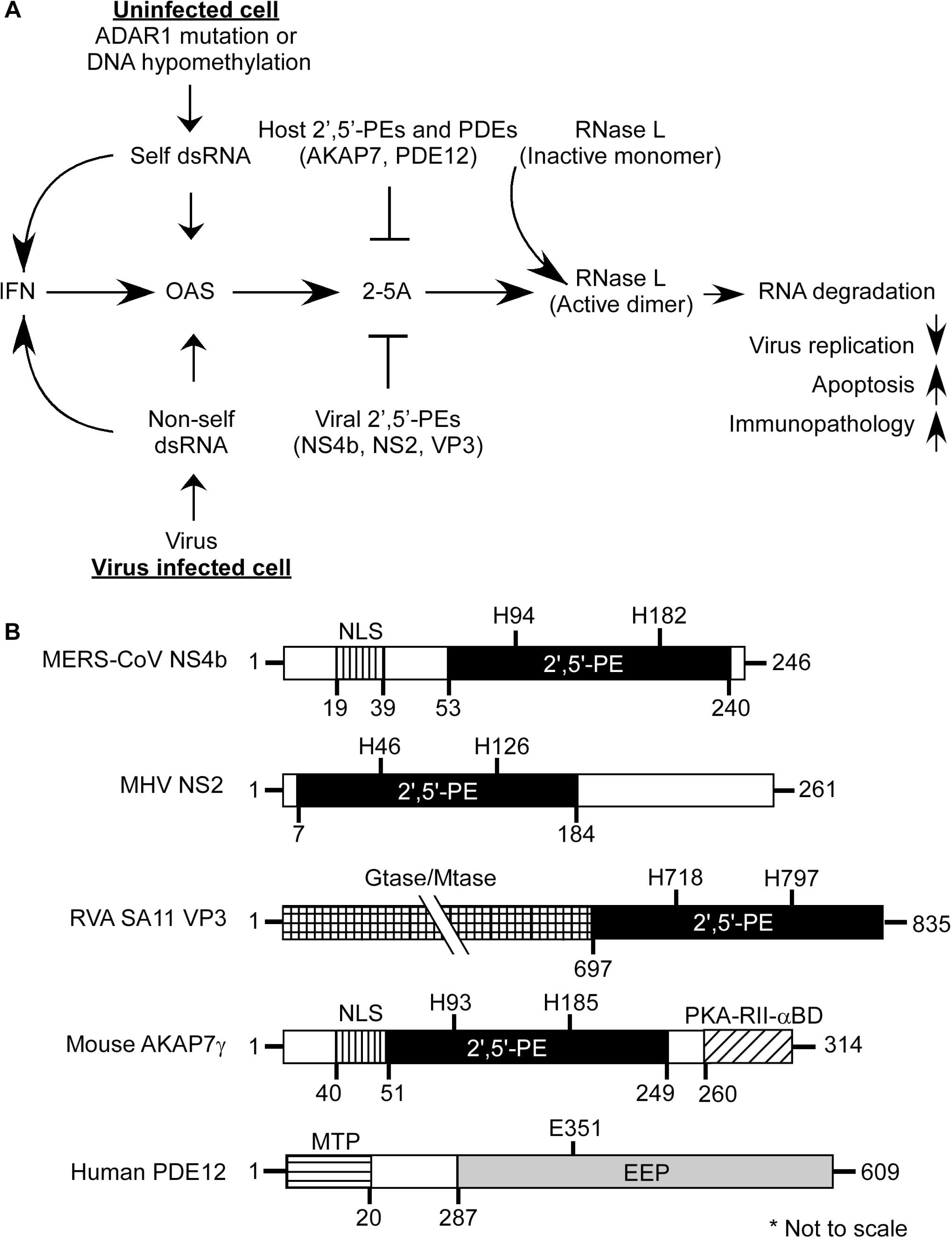
Interplay between cellular responses to viral and host dsRNA, the OAS-RNase L pathway, and antagonism by 2-5A degrading enzymes. (A) OASs(1–3) are IFN induced dsRNA sensors, once activated they synthesize the antiviral substance 2’,5’-oligoadenylates (2-5A) from ATP. 2-5A binds inactive monomeric RNase L inducing active RNase L dimers, which in turn degrade viral and host single-stranded RNAs. The balance between 2-5A accumulation by OAS enzymes and its degradation by host and viral enzymes determines cell and virus fate and inflammatory responses. (B) Domain structure of viral and cellular 2’,5’-PEs and human PDE12 (an endonuclease/exonuclease/phosphatase (EEP) family member). Features of full length MERS-CoV NS4b, MHV NS2, RVA SA11 VP3 and *M. musculus* AKAP7γ proteins including a nuclear localization sequence (NLS) and catalytic 2’,5’-PE domains are compared, modified from ref. (24). Position of conserved histidines within the catalytic domain of 2’,5’-PEs are shown. PKA-RII-α-BD, a binding domain for regulatory subunit (RII) of cAMP-dependent protein kinase A (27), guanylyltransferase (Gtase), and methyltransferase (Mtase) domains are also shown (25, 29). The mitochondrial-matrix targeting peptide (MTP) and the catalytic EEP domain of PDE12 is shown (55). Domains shown are not drawn to scale.

Regulation of 2-5A degradation is a key point of control in the OAS-RNase L pathway. Previously, we identified several different members of the eukaryotic-viral LigT group of the 2H-phosphoesterases (2H-PE) superfamily, named for two His-ɸ -Thr/Ser-ɸ motifs (where ɸ is a hydrophobic residue) that degrade 2-5A and therefore function as potent RNase L antagonists (19, 20) (Fig. 1B). Here we refer to 2H-PE members that degrade 2’,5’-oligoadenylates as 2’,5’-PEs. Other members of the 2H-PE superfamily have different activities, including 2’,3’-cyclic nucleotide phosphodiesterase and 3’,5’-deadenylase activities (19, 21).

The prototype of the 2’,5’-PEs is the mouse coronavirus (CoV), mouse hepatitis virus (MHV), accessory protein NS2 (22). However, predicted or confirmed 2’,5’-PEs are expressed by many betacoronaviruses (embecovirus lineage MHV, human coronavirus (HCoV) OC43, human enteric coronavirus (HECoV), equine coronavirus (ECoV), porcine hemagglutinating encephalomyelitis virus (PHEV) and merbeco lineage MERS-CoV and related bat CoVs), related toroviruses and group A and B rotaviruses (20, 23–26). However, the betacoronaviruses SARS-CoV and SARS-CoV-2 lack a 2’,5’-PE. Perhaps as a consequence, SARS-CoV-2 activates, and is inhibited by, the OAS and RNase L pathway (4). In addition, there is also a mammalian 2’,5’-PE, A-kinase anchoring protein (AKAP7, aka AKAP15 or AKAP18), that degrades 2-5A (27). Here we have expressed, purified and characterized the 2’,5’-PEs from MHV (NS2), MERS-CoV (NS4b), rotavirus group A (RVA) (VP3-C-terminal domain, CTD), and mouse AKAP7. We show that NS2 and NS4b are remarkably specific for cleaving 2’,5’-linked oligoadenylates, whereas AKAP7 and VP3-CTD will also cleave other 2’,5’- oligonucleotides. In contrast, all of the viral and mammalian 2’,5-PEs tested lack the ability to cleave 3’,5’-oligoribonucleotides. We further show that these enzymes are metal ion-independent and cleave trimer 2-5A (2’,5’-p_3_A_3_) producing mono- and di- adenylates with 2’,3’-cyclic phosphoryl termini. Our findings suggest that the sole function of the viral 2’,5’-PEs may be to eliminate 2-5A allowing some coronaviruses and rotaviruses to evade the antiviral activity of RNase L.

## RESULTS

### Phylogenetic relationship and alignment of viral and cellular 2’,5’-PEs

To probe the precise molecular mechanism by which 2’,5’-PEs allow some viruses to evade the antiviral effector RNase L, we further investigated MHV NS2, MERS-CoV NS4b, rotavirus group A (RVA) VP3-C-terminal domain (CTD), and mouse (mu)AKAP7, (Fig. 1B). A comparison of the domain organization of these 2’,5’-PEs shows a related, catalytic domain. Some of these enzymes have additional domains related to intracellular localization, nucleic acid metabolism or protein binding functions indicative of their cellular compartment-specific or accessory functions (Fig. 1B). For instance, MERS-CoV NS4b and muAKAP7 contain N-terminal nuclear localization signal (NLS) domains (24, 27, 28). VP3 is a multifunctional enzyme that contains N-terminal guanylyltransferase (Gtase) and methyltransferase (Mtase) domains involved in capping of the 5’ termini of viral mRNAs (25, 29). muAKAP7 also has a carboxy-terminal binding domain for the regulatory subunit II (RII) of cyclic AMP (cAMP)-dependent protein kinase A (PKA-RII-α-BD) (27). In addition, MHV NS2 protein has a C-terminal extension of unknown identity or function (Fig. 1B).

To determine the phylogenetic relationships between the different 2’,5’-PEs, we constructed a tree for amino acid sequences containing the catalytic domains from coronavirus, rotavirus and mammalian 2’,5’-PEs (Fig. 2A). 2’,5’-PEs were distributed into two distinct branches on the phylogenetic tree. VP3 group of proteins clustered into one branch while the other three groups containing NS2, NS4b and AKAP7 formed a separate branch (Fig. 2A). Within the VP3 group, RVA and RVB resolved on distinct sub-branches. Previously, full-length VP3 from RVA and RVB were also shown as separate distinct branches analogous to two separate clades (clade A and clade B) (30, 31). Interestingly, the NS2 proteins were mostly closely related to the mammalian AKAP7 catalytic domains, and then to the bat coronaviruses (HKU5 and SC2013) and MERS-CoV. The rotavirus VP3 proteins were most distally related to the other 2’,5’-PEs.

**Figure 2.**
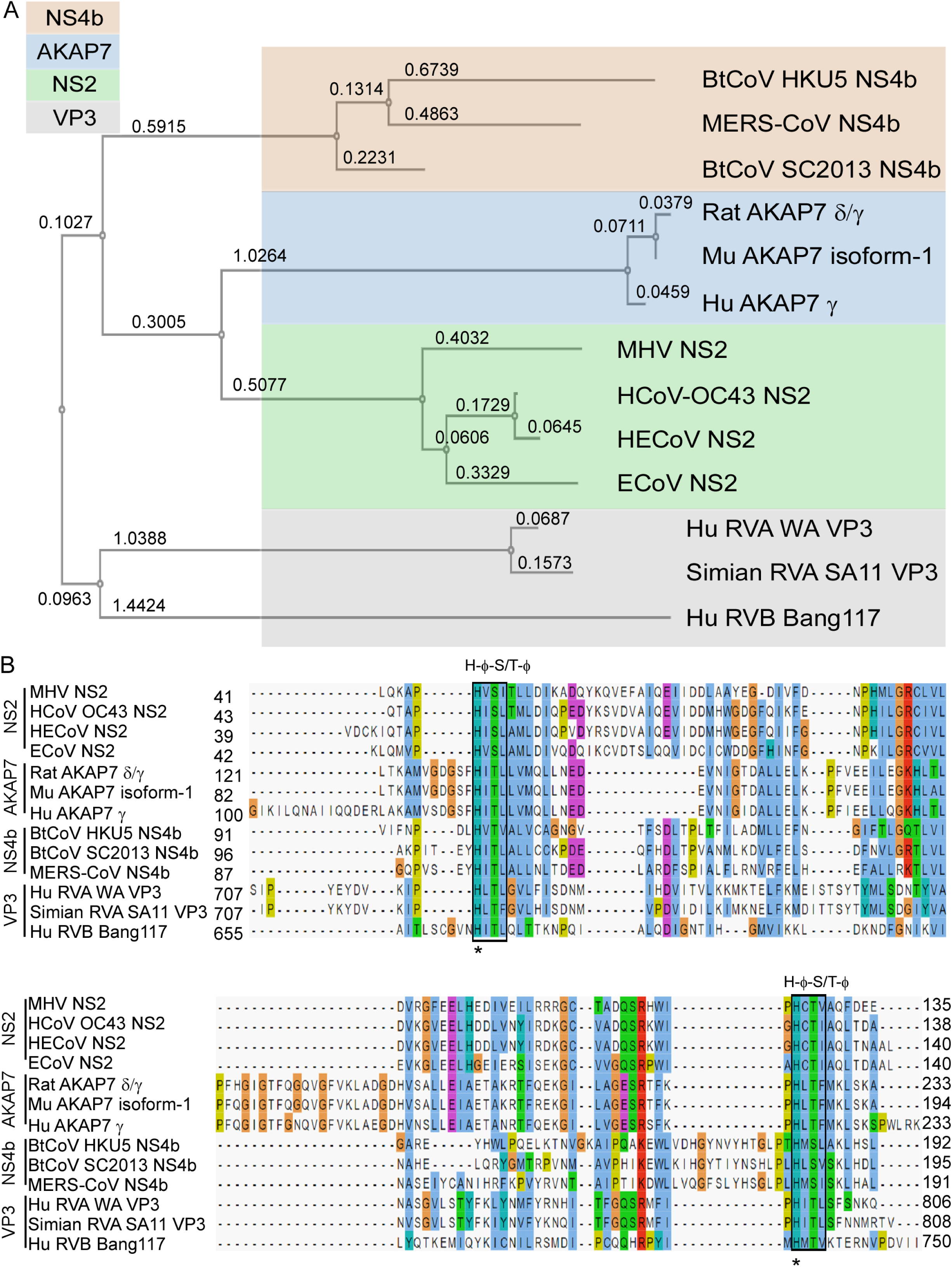
Phylogenetic relationship and sequence alignment of 2’,5’-PEs. **(A)** Phylogenetic tree based on amino acids from catalytic domains of 2’,5’-PEs is shown. The numbers represents branch length. (B) Sequence alignment of amino acids spanning the catalytic domain of 2’,5’-PEs using MAFFT multiple sequence alignment program is shown. Catalytic motifs [H-Φ-(S/T)-Φ] are indicated above the boxes, where Φ represents a hydrophobic residue. Numbers represent the start and end of the amino acid sequences used for sequence alignment. Aligned residues are color-coded for conservation according to the CLUSTAL X scheme. Hydrophobic, blue; positive charge, red; negative charge, magenta; polar, green; glycine, orange; proline, yellow; aromatic, cyan; unconserved, white. Hu, human; Mu, mouse; HE, human enteric; E, equine; Bt, bat; CoV, coronavirus; RVA, rotavirus group A; RVB, rotavirus group B.

Based on the phylogenetic relationship and functional relatedness, we further analyzed the sequence conservation by amino acid alignment. 2H-PE superfamily members are characterized by the presence of two H-ɸ-S/T-ɸ motifs, separated by an average of 80 amino acids (where ɸ represents a hydrophobic amino acid) (19). The alignment shows that both motifs are highly conserved across all 2’,5’-PE proteins (Fig. 2B, see boxes).

These motifs form the catalytic core that bind to and cleave the 2-5A substrate. Consistent with the phylogenetic analysis, sequence analysis revealed that 2’,5’-PEs clustered into four groups corresponding to NS2, NS4b, AKAP7 and VP3. The two histidines within the conserved motifs were 100% conserved among all the sequences (Fig. 2B, see asterisks). Several residues with intergroup consensus of >50% were identified in the alignment. The amino acid alignment shows several regions of conservation that exist beyond the two conserved catalytic motifs (H-ɸ-S/T-ɸ) (Fig. 2B, shown above sequence alignment).

Among the sequences in alignment, AKAP7 proteins of human, rat and mouse origin shared the highest amino acid identity ranging between 85 to 97% (88 to 97% similarity) (Table S1). NS2 proteins shared 48 to 92% identity (64 to 94% similarity), NS4b proteins shared 35 to 49% identity (50 to 69% similarity), and VP3 proteins shared 16 to 78% identity (29 to 84% similarity) within their groups. Interestingly while RVA VP3 proteins shared a high 78% identity (84% similarity) between them, they share only 16% identity (29% similarity) with the representative of RVB VP3 protein. The catalytic domains of 2’,5’-PEs have modest intragroup alignment and a relatively lower intergroup alignment. Overall intergroup alignment for the catalytic domains of these proteins shown 10 to 22% identity (19 to 36% similarity) (Table S1). NS2 proteins shared 11 to 16% amino acid identity (24 to 29% similarity) with NS4b, 16 to 22% identity (30-36% similarity) with AKAP7 and 16 to 22% identity (26 to 32% similarity) with VP3 proteins. Similarly, NS4b group of proteins shared 12 to 18% identity (21 to 29% similarity) with AKAP7 and 10 to 19% identity (23 to 34% similarity) with VP3 proteins. AKAP7 and VP3 shared 11 to 20% identity and 19 to 25% similarity between the two groups.

### 2’,5’-PEs are specific for 2’,5’-linked phosphodiester bonds and preferably cleave 2’,5’-oligoadenylate

2’,5’-PEs are members of the LigT family of the 2H-PE superfamily of enzymes, which are involved in RNA processing that can act on diverse substrates (19). Also, members such as MERS-CoV NS4b and muAKAP7 have a functional NLS peptide (24, 27). To determine if there was a wider role for these enzymes beyond cleaving 2-5A, we tested an expanded set of potential substrates. MERS-NS4b, MHV NS2, RVA VP3-CTD and muAKAP7 proteins were expressed in bacteria and then purified. Also, for comparison, human PDE12 (aka 2’-PDE), a member of the exonuclease-endonuclease-phosphatase (EEP) family known to cleave 2-5A was purified (Fig. 1B) (32, 33). The catalytically inactive mutant proteins, MERS-NS4b^H182R^, MHV NS2^H126R^, RVA VP3-CTD^H718A^, muAKAP7^H93A; H185R^ and human PDE12^E351A^ served as the negative controls. Purity and identity of trimer 2-5A (2’,5’-p_3_A_3_) were confirmed by HPLC (Fig. 3A) and mass spectrometry [described later for (Fig. 5J)]. Purified 2’,5,-PE proteins were incubated with 2-5A substrate at 30°C for 1 h and the 2-5A cleavage products analyzed by HPLC using a C18 column. All five wild type proteins cleaved 2-5A as observed by loss of intact 2-5A and appearance of peaks for the different cleavage products (Fig. 3B-F). Interestingly, MERS-NS4b (Fig. 3B) and MHV NS2 (Fig. 3C) degraded 2-5A to give two prominent products whereas RVA VP3-CTD (Fig. 3D) and muAKAP7 (Fig. 3E) gave four products upon extended degradation of 2-5A suggesting a difference in either mechanism or rate of cleavage by these proteins. On the other hand, 2-5A cleavage by human PDE12 (Fig. 3F) results in the formation of two products corresponding to the elution time of the standard ATP and 5’-AMP, as previously described (32). As expected, the 2’,5’-PE catalytically inactive mutant proteins containing a His-to-Arg or His-to-Ala mutations in the conserved histidines did not cleave 2-5A (Fig. 3G-J). Human PDE12 with Glu-to-Ala mutation at 351 amino acid residue also did not degrade 2-5A, as described previously (34) (Fig. 3K). Importantly, these findings show a different mode of 2-5A cleavage between 2’,5’-PEs, members of the 2H-PE superfamily, and PDE12, a member of the EEP family of phosphodiesterases.

**Figure 3.**
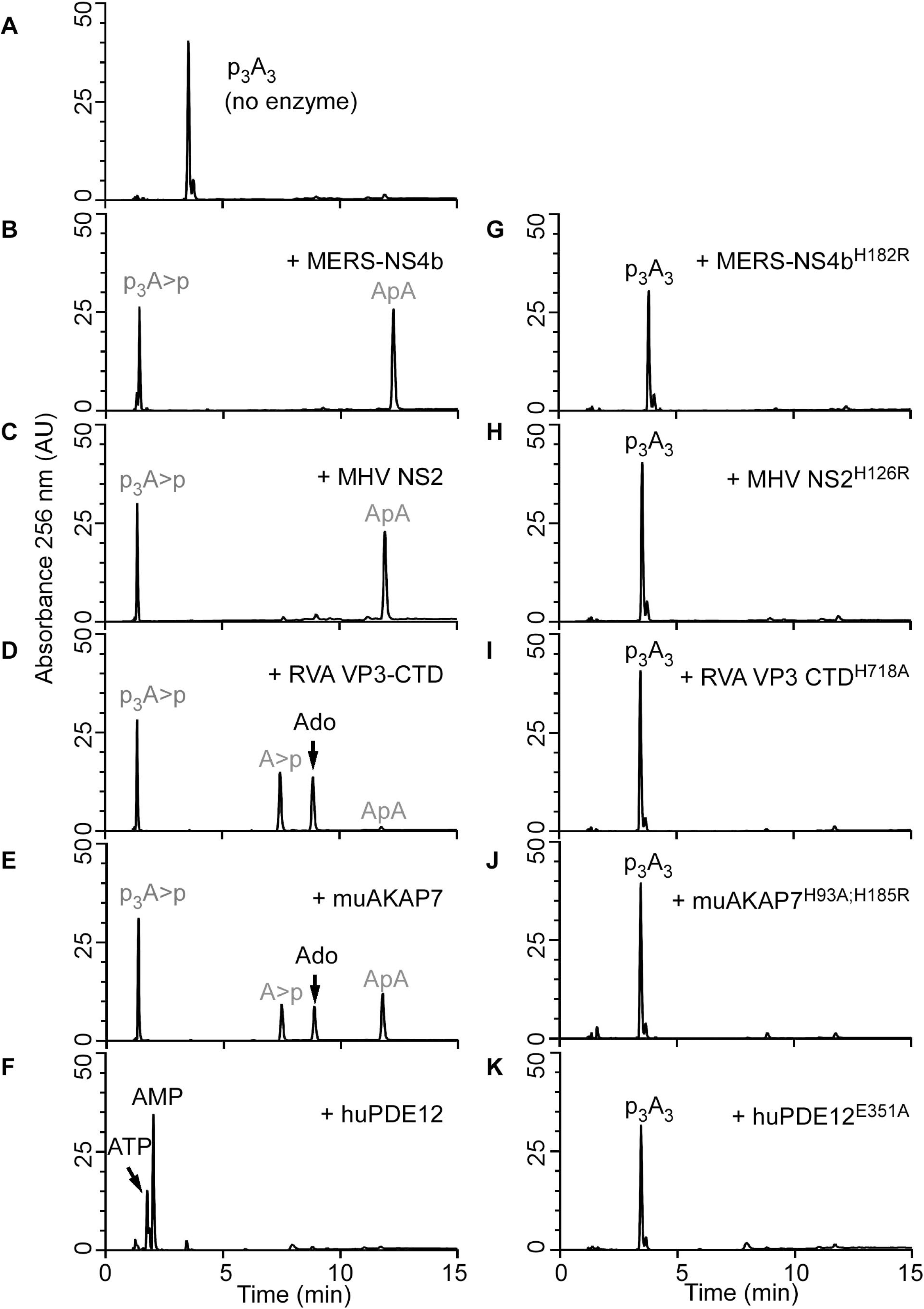
Specific cleavage of trimer 2-5A (2’,5’-p_3_A_3_) by viral and cellular 2’,5’- PEs. HPLC analysis of (A) intact 2’,5’-p_3_A_3_, followed by its cleavage with either viral 2’,5’-PEs (B) MERS-NS4b, (C) MHV NS2, (D) RVA VP3-CTD or a host 2’,5’-PE (E) muAKAP7 or (F) human PDE12. Purified 2’,5’-p_3_A_3_ (200 µM) was incubated with 1µM of indicated proteins at 30°C for 1 h. HPLC analysis of catalytically inactive mutants of these enzymes incubated with 2’,5’-p_3_A_3_ under similar conditions is shown for (G) MERS-NS4b^H182R^ (H) MHV NS2^H126R^ (I) RVA VP3-CTD^H718A^ (J) muAKAP7^H93A; H185R^ and (K) human PDE12^E351A^. Experiments performed at least three times produced a similar 2’,5’-p_3_A_3_ degradation pattern for each 2’,5’-PEs. Arrows indicate elution times of the standards ATP, AMP, and adenosine (Ado). Peaks shown in grey were determined from experiments done in figures 5 and 6.

To investigate the expanded substrate specificity of 2’,5’-PEs, we tested possible cleavage of various 2’-5’ and 3’-5’ linked oligoribonucleotides by HPLC. Purified 2’,5’-PE proteins (1 µM) were incubated with either 2’-5’- or 3’-5’-linked pentaribonucleotide substrates (10 µM) at 30^°^C for 1 h. Wild type MERS-NS4b, specifically degraded 2’-5’ p5’(rA)_5_ by >99% while 2’-5’ p5’(rU)_5_, p5’(C)_5_ or p5’(G)_5_ were not degraded (≤4%) (Table S2). Catalytically inactive mutant MERS-NS4b^H182R^ did not degrade any of the tested substrates under similar conditions. Wild type MHV NS2 also specifically degraded 2’-5’ p5’(rA)_5_ >99% while 2’-5’ p5’(rU)_5_, p5’(C)_5_ or p5’(G)_5_ were not degraded (≤7%) (Table S2). Mutant MHV NS2^H126R^ did not degrade any of the tested substrates. These results suggest MERS-NS4b and MHV NS2 are remarkably specific in degrading 2’-5’ linked oligoadenylate compared to the other substrates. We further tested RVA VP3-CTD which degraded 2’-5’ p5’(rA)_5_ >95%, p5’(rU)_5_ ∼ 40%, p5’(C)_5_ ∼90%, and p5’(G)_5_ ∼6% while mutant RVA VP3-CTD^H718A^ did not degrade any of the tested substrates (Table S2). Wild type muAKAP7 degraded 2’-5’ p5’(rA)_5_ >99%, p5’(rU)_5_ >95%, p5’(C)_5_ >95%, and p5’(G)_5_ >90% while mutant muAKAP7^H93A;H185R^ did not degrade any of the tested substrates with the exception of 2’-5’ p5’(G)_5_ ∼40% (Table S2). To ensure that exclusive cleavage of 2’,5’-oligoadenylates by MERS-NS4b was not due to limiting amounts of enzyme, 10 µM of different 2’,5’-linked penta-ribonucleotides were incubated with three-fold higher concentrations (3 µM) of MERS-NS4b at 30°C for 1h. Wild type MERS-NS4b specifically degraded 2’-5’ p5’(rA)_5_ >99% while 2’-5’ p5’(rU)_5_, p5’(C)_5_ or p5’(G)_5_ were not degraded (≤6%) suggesting MERS-NS4b enzymatic activity is specific for degradation of 2’,5’-oligoadenylates (Table S3). Because MERS NS4b and MHV NS2 cleaved 2’-5’ p5’(rA)_5_ but not other 2’-5’-linked substrates, we further determined if this was due to lack of binding to the other substrates. To test this possibility, 10 µM of 2’-5’ p5’(rA)_5_ was incubated with 0.2 µM of MHV NS2 in the absence or presence of increasing concentrations of 2’-5’ p5’(rU)_5_ at 30°C for 10 min. The amounts of 2’-5’ p5’(rA)_5_ degraded by MHV NS2 in the presence of 0, 3.1, 10, 12.5, 25, 50 and 100 µM was determined by HPLC analysis (Fig. S1). Degradation of 2’-5’ p5’(rA)_5_ by MHV NS2 decreased as the amount of 2’-5’ p5’(rU)_5_ in the reaction increased beyond 10 µM (i.e. ratio >1:1) (Fig. S1). Our results suggests that 2’-5’ p5’(rU)_5_ was able to bind MHV NS2 and competitively interfere with MHV NS2 ability to cleave 2’-5’ p5’(rA)_5_.

We next tested degradation activity of 2’,5’-PEs against 3’-5’ linked p5’(rA)_5_, p5’(rU)_5_ and p5’(C)_5_. One µM of enzyme was incubated with 10 µM of the substrate at 30°C for 1 h. Wild type MERS-NS4b and its mutant MERS-NS4b^H182R^, wild type MHV NS2 and its mutant NS2^H126R^, RVA VP3-CTD and its mutant RVA VP3-CTD^H718A^, and wild type muAKAP7 and its mutant muAKAP7^H93A; H185R^ (Table S2) did not degrade the 3’-5’ linked substrates 3’-5’ p5’(A)_5_, 3’-5’ p5’(U)_5_, and 3’-5’ p5’(C)_5_. [We were unable to obtain 3’-5’ p5’(G)_5_ because of repeated failures of its chemical synthesis and/or purification, therefore this oligonucleotide could not be tested]. Our results suggest that all of the 2’,5’-PEs examined are highly specific for cleaving 2’,5’ linked oligoribonucleotides. Among 2’,5’ linked substrates MERS-NS4b and MHV NS2, are specific for cleaving 2’-5’ oligoadenylate, whereas RVA VP3-CTD cleaved in order: 2’-5’ pA5>pC5>pU5>>pG5 and muAKAP7 cleaved all of 2’,5’ linked pentanucleotides with similar efficacy.

Based on the differential enzymatic activity of these 2’,5’-PEs in degrading different types of 2’,5’-linked phosphodiester substrates, we tested if they could degrade 2’,3’-cyclic-GMP-AMP (cGAMP). cGAMP is a cyclic-dinucleotide secondary messenger with mixed phosphodiester linkages between 2’-OH of GMP to 5’-phosphate of AMP and 3’-OH of AMP to 5’-phosphate of GMP, synthesized by cyclic GMP-AMP-synthase (cGAS) in response to cytoplasmic dsDNA (35). cGAMP was incubated either with or without wild type and mutant 2’,5’-PEs at 30°C for 1h and analyzed by HPLC. Wild type MERS-NS4b, MHV NS2, RVA VP3-CTD, and muAKAP7 did not degrade 2’,3’-cGAMP whereas they did degrade 2’,5’-p_3_A_3_ (served as a positive control) under similar conditions (Table S4). Catalytic mutants of 2’,5’-PEs tested did not degrade 2’,3’- cGAMP or 2’,5’-p_3_A_3_ under similar conditions. The results suggest that 2’,5’-PEs are capable of cleaving 2’,5’-phosphodiester bonds in linear homo-ribonucleotides but not in the cyclic-mixed phosphodiester linked 2’,3’-cGAMP.

### 2’,5’-PEs exhibit metal ion-independent phosphodiesterase activity

Metal ion dependency was evaluated by performing assays in either the presence of EDTA or magnesium. In the presence of EDTA, without added magnesium, 1 µM of MERS-NS4b (Fig. 4A) or MHV NS2 (Fig. 4B) degraded >90% of 2-5A in ∼20 min whereas 0.05 µM of RVA VP3 CTD (Fig. 4C) and 1µM of muAKAP7 (Fig. 4D) degraded >90% of the 2-5A within 5 min. Relative rates of 2-5A degradation by RVA VP3-CTD >> muAKAP7 > MHV NS2 = MERS-NS4b were observed. Based on the specific activities, the ratio of fold activity of RVA VP3-CTD: muAKAP7: MERS-NS4b: MHV NS2 was 38.9: 2.9: 1: 1. It is noteworthy that many mammalian cell types have a total cellular Mg^2+^ concentration in between 17 to 20 mM, of which only 5-22% may be free depending on the cellular compartment. (36). We determined the specific activities of the 2’,5’-PEs for degrading 2-5A in the absence and presence of 10 mM MgCl_2_ at 5 min (Fig. 4E). The addition of Mg^2+^ ions decreased the specific activity of MERS-NS4b to ∼0.6 fold and that of muAKAP7 to ∼0.8 fold. The activity of MHV NS2 and RVA VP3-CTD showed a negligible decrease to 0.97 fold and 0.99 fold in the presence of Mg^2+^ ions, respectively. Our results suggest that the 2’,5’-PEs activity of these proteins is independent of Mg^2+^ ions and its presence either slightly decreases or has no effect on the activity of these enzymes.

**Figure 4.**
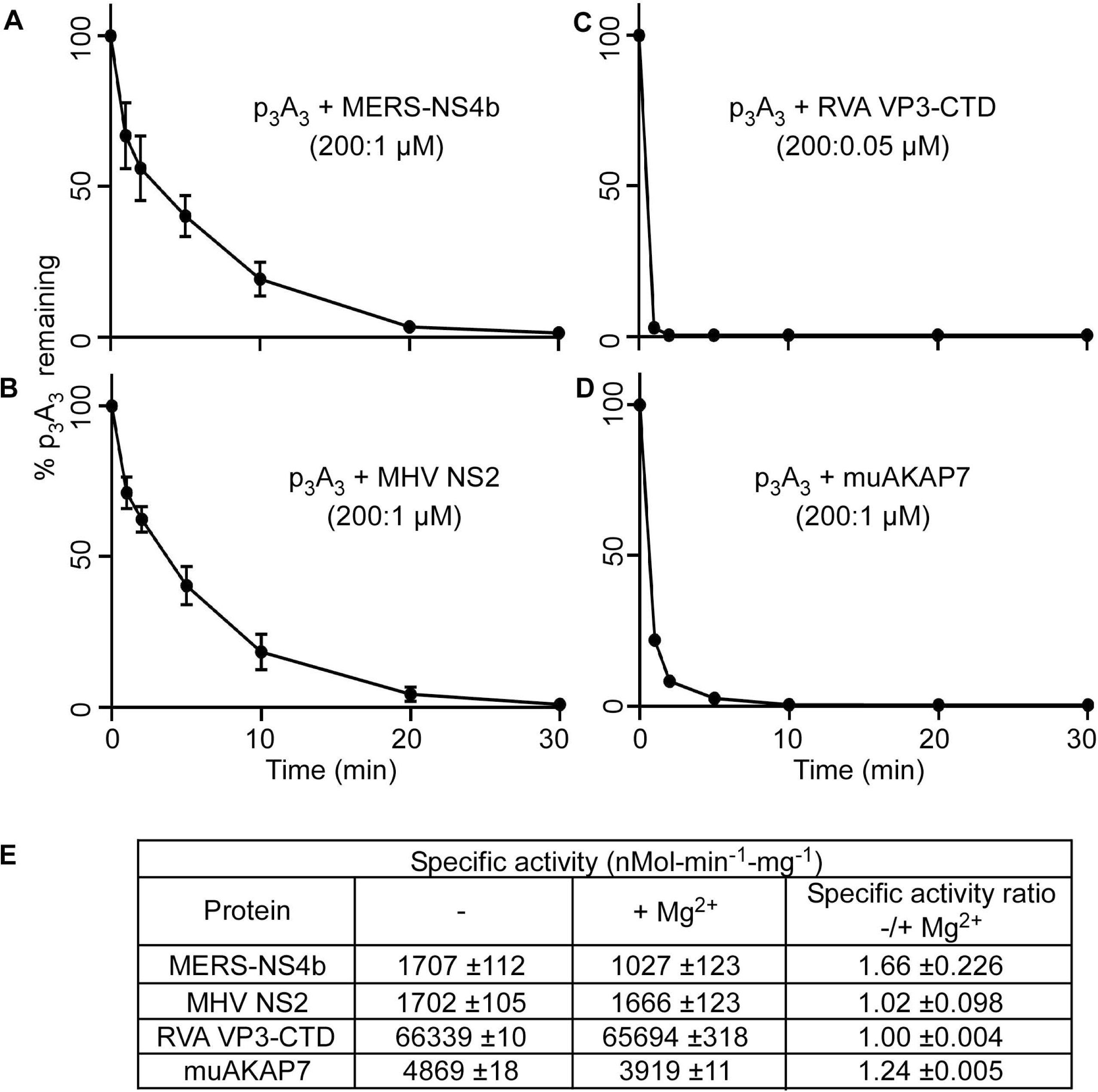
Influence of Mg^2+^ ions on degradation of 2’,5’-p_3_A_3_ by 2’,5’-PEs. Purified 2’,5’-p_3_A_3_ was incubated with indicated 2’,5’-PEs in time-course experiments. Data was obtained by incubating 2’,5’-p_3_A_3_ (200 µM) with (A) MERS-NS4b (1µM), (B) MHV NS2 (1 µM), (C) RVA VP3-CTD (0.05 µM) and (D) muAKAP7 (1 µM) at 30°C. Samples were collected at 1, 2, 5, 10, 20 and 30 min and reactions were stopped. The percent of uncleaved 2’,5’-p_3_A_3_ remaining at indicated times were determined by calculating the area under the peaks on the HPLC chromatograms. (E) The table shows the specific activity of 2’,5’-PEs in the absence and presence of 10 mM MgCl_2_. Activity is expressed as the amount of products released from the substrate in nMol per min per mg of the protein at 30°C during 5 min reaction time. Experiments were performed in triplicate (n = 3) and the standard error of mean was calculated.

### 2’,5’-PEs cleave 2’,5’-linked oligoadenylate leaving products with cyclic 2’,3’ phosphoryl termini

Differences in the 2-5A cleavage products as determined by HPLC (Fig. 3) suggested that viral and mammalian 2’,5’-PEs cleave 2-5A via a different mechanism than human PDE12, which degrades 2-5A to produce ATP and AMP (32). Among 2’,5’-PEs, MERS-NS4b and MHV NS2 degraded 2-5A to give two cleavage products whereas RVA VP3-CTD and muAKAP7 gave four cleavage products. Therefore we decided to determine the precise cleavage sites in 2-5A by 2’,5’-PEs. 2-5A was partially digested with MERS-NS4b (Fig. 5) or RVA VP3-CTD (Fig. 6) followed by the collection of individual peak fractions of cleavage products. Cleavage products were subsequently identified and confirmed by HPLC analysis (comparing elution time with known standards), identification by m/z ratio (LC/MS/MS analysis of collected peaks) or biochemical analysis (by 5’-dephosphorylation) (Figs. 5A & 6A).

**Figure 5.**
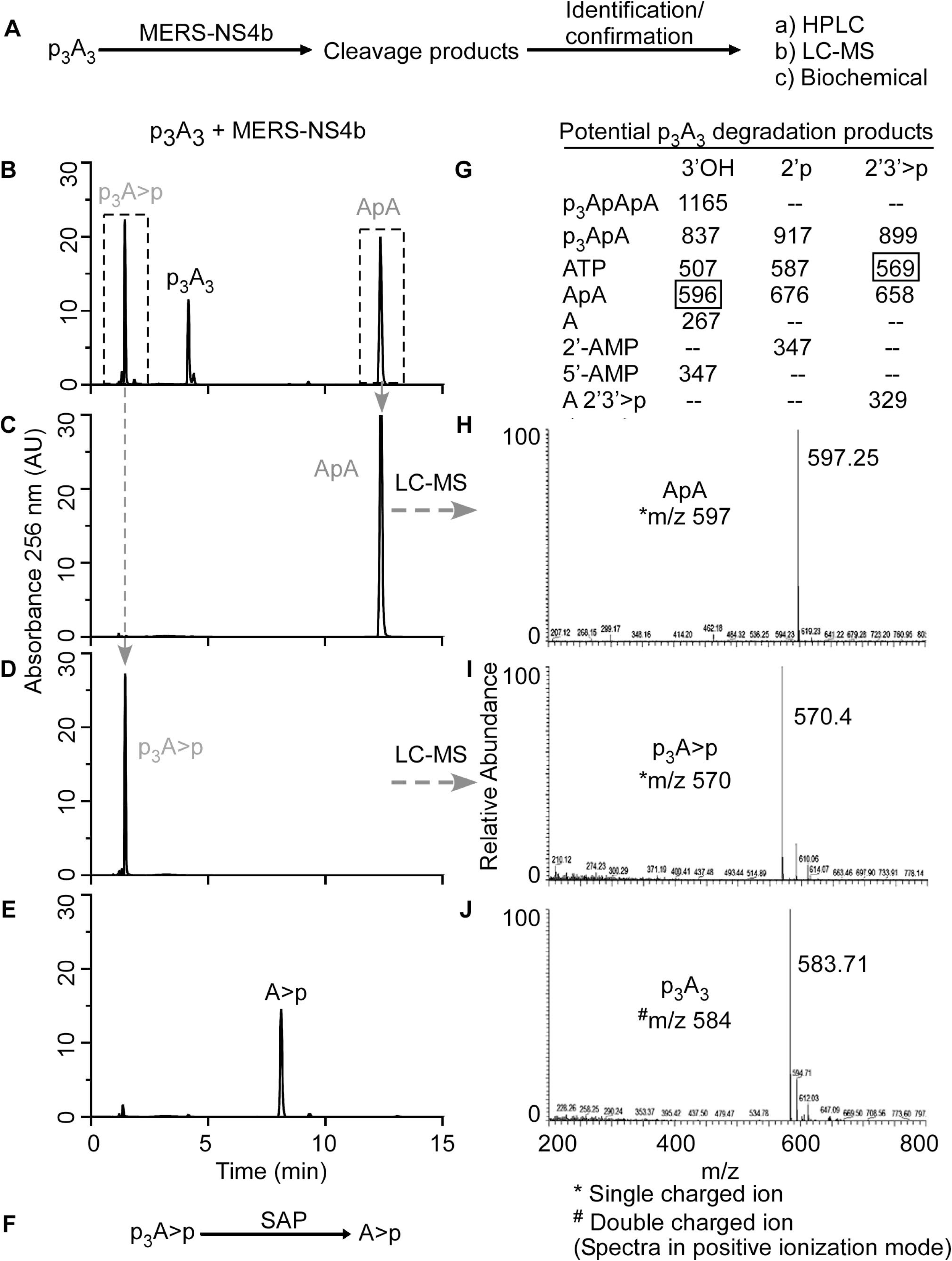
MERS-NS4b cleaves 2’,5’-p_3_A_3_ and catalyzes the formation of 2’,3’ cyclic phosphate products. (A) Schematic illustration of the strategy to identify the cleavage mechanism and degradation products of 2’,5’-p_3_A_3_ by MERS-NS4b. (B) Chromatogram of partially degraded 2’,5’-p_3_A_3_ and cleavage products formed by MERS-NS4b. 200 µM of 2’,5’-p_3_A_3_ was incubated with 1 µM of MERS-NS4b at 30°C for 10 min. (C) HPLC chromatogram of the collected peak (corresponds to ApA). (D) HPLC chromatogram of the collected peak (corresponds to p_3_A>p). (E) HPLC analysis of the dephosphorylated product (A>p) of peak collected in figure 5D (p_3_A>p). (F) Schematic illustration showing shrimp alkaline phosphatase (SAP) mediated p_3_A>p dephosphorylation at 5’ end forms A>p. (G) Expected masses of potential 2’,5’-p_3_A_3_ degradation products containing 3’OH, 2’p or 2’,3’>p groups. Box shows masses of actual cleavage products identified by mass spectrometry. Mass spectrometry analysis showing m/z of (H) ApA ◊ peak fraction collected in C, (I) p_3_A>p ◊ peak collected in D and (J) intact 2’,5’-p_3_A_3_. m/z is the mass-charge ratio. Peaks shown in grey were identified in the subsequent experiments.

**Figure 6.**
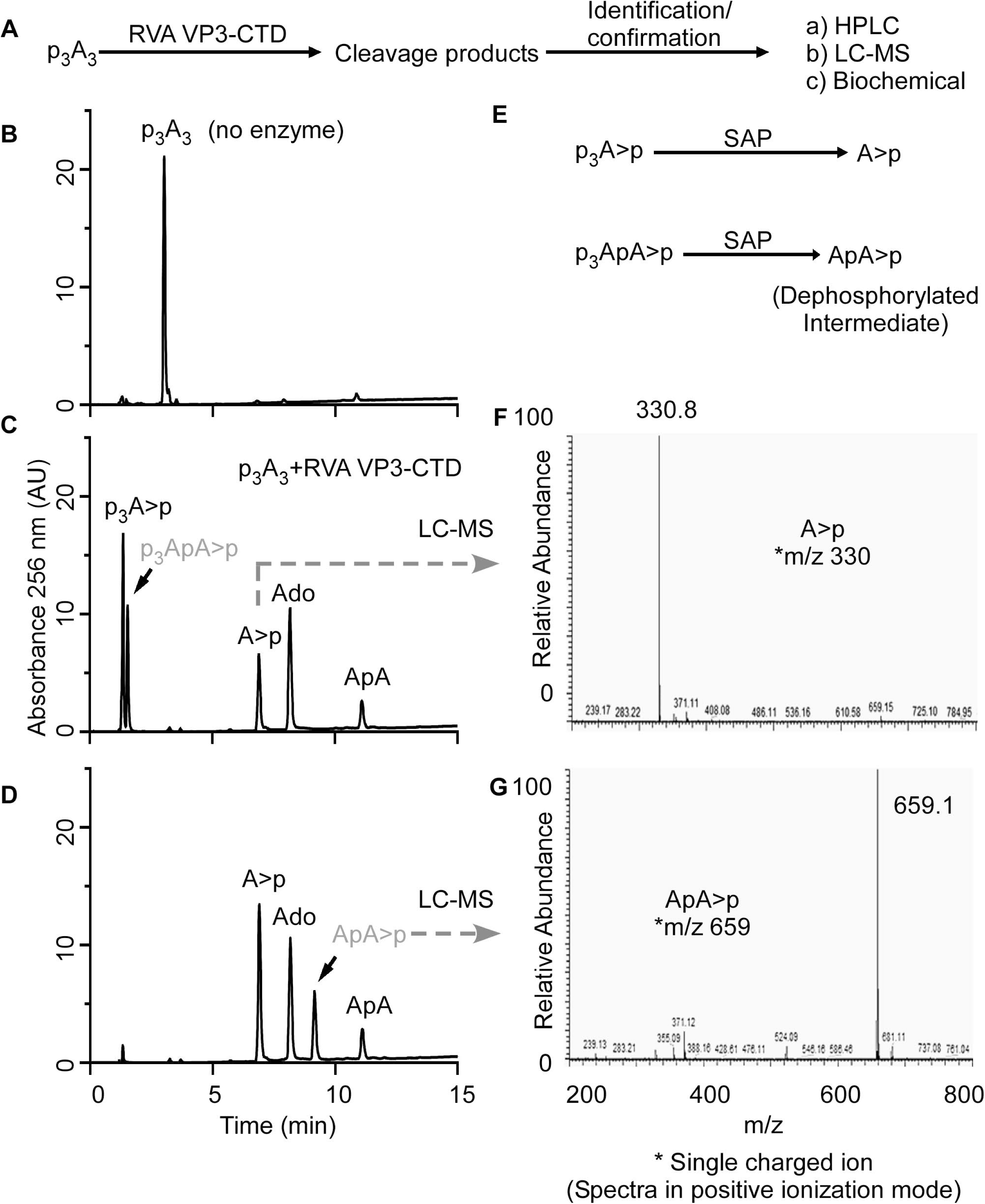
2’,5’-p_3_A_3_ cleavage by RVA VP3-CTD proceeds via formation of p_3_ApA>p and ApA intermediates. (A) Schematic illustration of the strategy to identify the cleavage mechanism, intermediates and end products of 2’,5’-p_3_A_3_ cleavage by RVA VP3-CTD. (B) Chromatogram of intact p_3_A_3_. (C) HPLC chromatogram of cleavage products formed by degradation of 200 µM 2’,5’-p_3_A_3_ incubated with 0.05 µM of RVA VP3-CTD for 20 min at 30°C. Peaks identified based on elution times of known standards are marked. (D) HPLC analysis of the dephosphorylated products from the reaction in figure 6C. Shrimp alkaline phosphatase (SAP) treatment dephosphorylate 5’- phosphates. The peaks indicated were collected and identified by LC/MS/MS analysis. (E) Schematic illustration showing dephosphorylation of potential intermediates at 5’ end using SAP. Mass spectrometry analysis showing m/z of (F) A>p and (G) ApA>p. m/z is the mass-charge ratio. Peaks shown in grey were identified in the subsequent experiments.

Partial digestion of 2-5A by MERS-NS4b was confirmed by HPLC analysis of the samples using a C18 column (Fig. 5B) and comparing it with the chromatogram of intact 2-5A (Fig. 3A). The partially digested 2-5A by MERS-NS4b was run on a Dionex DNAPac^R^PA-100 column in an ammonium bicarbonate volatile buffer system as described in Methods. Individual peaks were collected and processed for mass spectrometry analysis. Individually collected peaks were also rerun on a C18 column to confirm the purity and matched elution times of the collected peaks before performing LC/MS/MS (Figs. 5C, D). Mass spectrometric analysis of the peaks revealed an m/z ratio of 597.25 (Fig. 5H) and 570.4 (Fig. 5I). The m/z ratios of 597.25 and 570.4 were compared to the masses of potential 2-5A degradation/intermediate products (Fig. 5G) and were found to correspond to ApA and p_3_A>p (where “>p” represents a 2’,3’ cyclic phosphate), respectively. Intact 2-5A gave an m/z ratio of 584 for the double charged ion (Fig. 5J). Moreover, the collected peak of p_3_A>p (shown in Fig. 5D) was subjected to SAP mediated 5’ dephosphorylation which results in the peak corresponding to A>p (Fig. 5E, F). This experiment suggested that MERS-NS4b degrades 2-5A to produce a 5’-product with 2’,3’ cyclic phosphate terminus in the form of p_3_A>p and a 3’-product of ApA. To test if p_3_A>p and ApA are end products of the reaction, we subjected 2-5A to extended degradation by MERS-NS4b and monitored the area under the peak corresponding to ApA at 0 h, 1 h, 4 h and 24 h (Fig. S2 A-D, I). After the ApA peak appears (at 1h) its amount remained unchanged up to 24 h. Also, 2’-5’ linked 5’pApA incubated with MERS-NS4b did not result in any degradation (Fig. S2 E-H, I). These results suggest that MERS-NS4b does not cleave di-adenylates into smaller products irrespective of 5’ mono-phosphorylation status under the given experimental conditions.

With a similar approach, we designed an experiment to elucidate the cleavage products and intermediates formed upon 2-5A degradation by RVA VP3-CTD (Fig. 6A). Because RVA VP3-CTD has high specific activity against 2-5A (Fig. 4), 2-5A was incubated with decreased protein concentrations and times of incubation to capture any possible intermediates and degradation products of 2-5A (Fig. 6B, C). 2-5A cleaved by VP3-CTD forms products which based on the elution times of the known standards and compounds were identified as p_3_A>p, A>p, ApA, adenosine and an unknown intermediate (shown in grey color) (Fig. 6C). Based on potential degradation intermediates we speculated the unknown intermediate to be p_3_ApA>p. To test this possibility, a part of the sample reaction with 2-5A cleavage products (obtained from the sample used in Fig. 6C) was subsequently treated with SAP to remove 5’- phosphorylation from cleavage products (if any) which would result in the formation of A>p and ApA>p from p_3_A>p and p_3_ApA>p, respectively (Fig. 6E). The dephosphorylated sample was analyzed by running it on a C18 column. After SAP treatment, the amount of adenosine and ApA remained constant when compared before (Fig. 6C) and after (Fig. 6D) dephosphorylation, as calculated by integrating the area under the peaks of the HPLC chromatograms. However, the total area under the peak corresponding to A>p increased suggesting A>p was formed as a result of dephosphorylation of p_3_A>p (Fig. 6C, D). Importantly, a new peak (possibly ApA>p) appears which is formed by dephosphorylation of an unknown intermediate (Fig. 6D, shown in grey color). The dephosphorylated samples were run on a Dionex DNAPac^R^ PA-100 column in an ammonium bicarbonate volatile buffer system as described in the Methods. Individual peak fractions were collected and processed for mass spectrometry analysis. Collected peaks were rerun on a C18 column to confirm the purity and match the elution time of the collected peak with that of A>p and ‘dephosphorylated intermediate’ (Fig. 6D) before performing LC/MS/MS. Mass spectrometric analysis of the peaks revealed m/z ratios of 330 corresponding to A>p (Fig. 6F) and 659.1 which corresponds to that of ApA>p (Fig. 6G). This experiment suggests that trimer 2-5A (2’,5’p_3_A_3_) degradation by RVA VP3- CTD proceeds via formation of p_3_ApA>p and ApA intermediates. RVA VP3-CTD degrades p_3_ApA>p to form p_3_A>p (5’-product) and A>p (3’-product) whereas di-adenylate (ApA) is further degraded to yield A>p (5’-product) and adenosine (3’- product). Complete degradation of p_3_A_3_ by RVA VP3-CTD results in the formation of p_3_A>p, A>p and adenosine as end products (Fig. 3D). Further, the preferred site of p_3_A_3_ cleavage by RVA VP3-CTD was investigated in a time-course experiment. RVA VP3-CTD (0.05 µM) was incubated with p_3_A_3_ (200 µM) substrate at 30°C and samples were collected at different time points. The substrate or product peaks at each time point were analyzed by calculating the percent of the area under the peaks of the HPLC chromatograms (Fig. S3) and tabulated (Fig. S4A). The analysis revealed that the majority of p_3_A_3_ is cleaved by RVA VP3-CTD to produce p_3_ApA>p (5’-product) and adenosine (3’-product). The intermediate species (p_3_ApA>p) is subsequently cleaved to produce p_3_A>p (5’-product) and A>p (3’-product). A minor fraction of p_3_A_3_ is cleaved by RVA VP3-CTD to produce p_3_A>p (5’-product) and ApA (3’-product). The di-adenylate intermediate (ApA) is subsequently cleaved into A>p (5’-product) and adenosine (3’- product) (Fig. S4B), which is apparent from incubations at a higher concentration of RVA VP3-CTD (Fig. 3D). Moreover, the degradation pattern of the two di-adenylate intermediates reveals that tri-phosphorylated intermediate (p_3_ApA>p) is readily cleaved by RVA VP3-CTD whereas ApA cleavage is slow suggesting 5’-tri-phosphorylated molecules are preferred over non-phosphorylated substrates (Figs. S3, S4). In a separate experiment, 10 µM of 2’-5’ linked 5’p(A)_2_ was incubated with 1 µM of VP3-CTD at 30°C for 1 h. The results confirmed the formation of cleavage products corresponding pA>p (5’-product) and adenosine (3’-product) (Fig. S5). Moreover, in another time-course experiment muAKAP7 cleaved p_3_A_3_ to produce p_3_A>p (5’-product) and ApA (3’-product) (Fig. S6). The ApA intermediate was further cleaved to form A>p and adenosine. Interestingly, unlike RVA VP3-CTD, muAKAP7 mediated cleavage of p_3_A_3_ does not form p_3_ApA>p intermediate.

Overall mechanisms and differences in degradation of 2-5A by representative EEP (PDE12) and 2’,5’-PEs family members are summarized in figure 7. Human PDE12 degrades trimer 2-5A into ATP and 2 (5’-AMP)s in the presence of Mg^2+^ ions, as has been reported (32). On the other hand, mammalian and viral 2’,5’-PEs, act in a metal-ion independent way, degrading 2-5A to form 5’ products with 2’,3’ cyclic phosphates. All 2’,5’-PEs quickly cleave active anti-viral 2-5A into inactive molecules that is, the products are not capable of activating RNase L because of a requirement for at least three adenylyl residues (37). MERS-NS4b and MHV NS2 degrade trimer 2-5A to form p_3_A>p and ApA. RVA VP3-CTD and muAKAP7 further cleave ApA to form A>p and adenosine as products. In addition to the above-mentioned degradation intermediates and products of 2-5A, RVA VP3-CTD also produced p_3_ApA>p as an intermediate, suggesting it to be a 2’,5’-specific endoribonucleolytic phosphodiesterase (Figs. 6 & 7).

**Figure 7.**
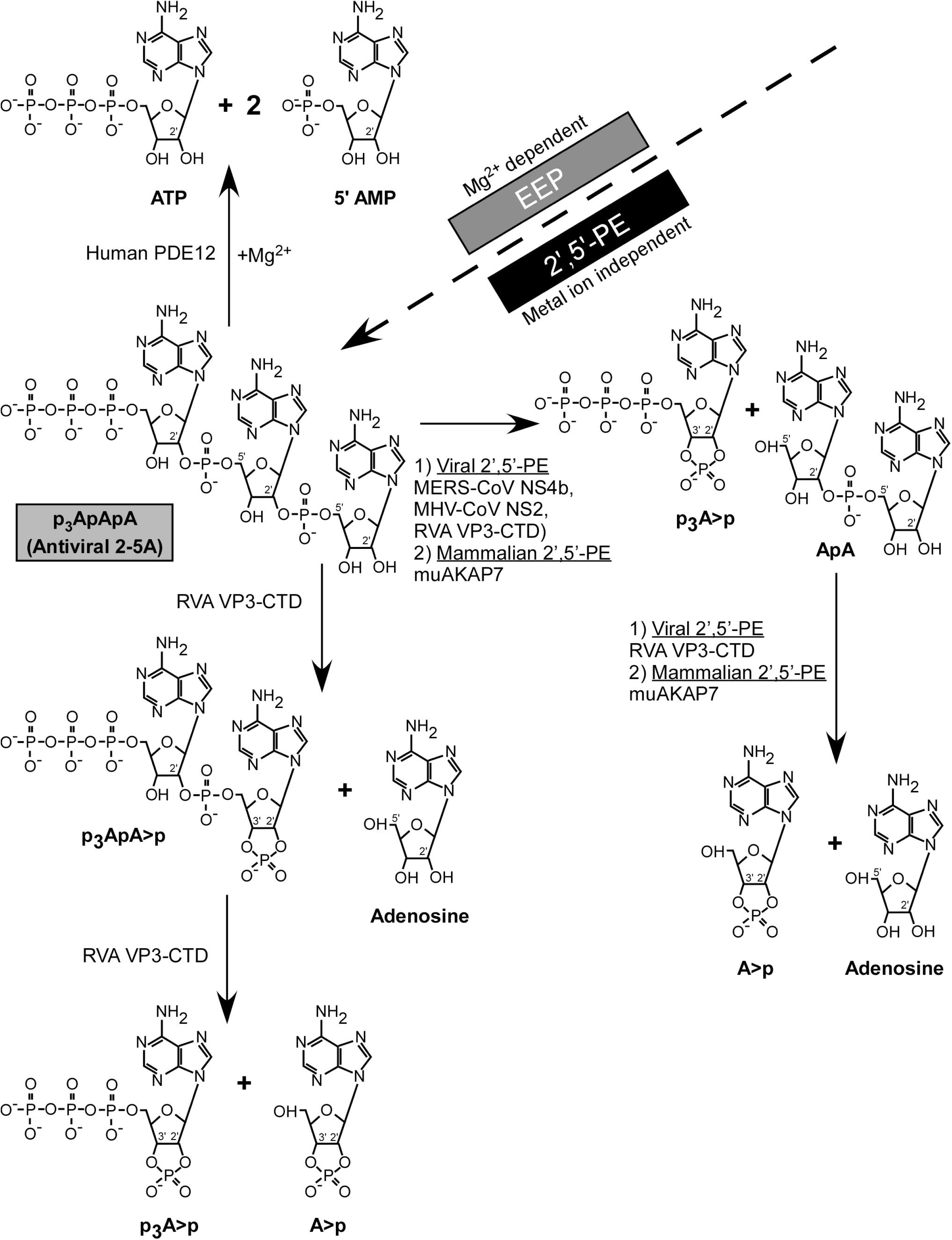
Mechanism of 2’-5’-p_3_A_3_ degradation by 2’,5’-PEs (a subfamily of 2H-PEs superfamily) and EEP (endonuclease/exonuclease/phosphatase) family. MERS-CoV NS4b, MHV-CoV NS2, RVA VP3, and mammalian mouse AKAP7 from the 2’,5’-PE subfamily cleave 2’,5’-p_3_A_3_ and leaves 2’,3’ >p groups on the 5’-products. While human PDE12, an EEP family member degrades 2’,5’-p_3_A_3_ to yield ATP and AMP.

## DISCUSSION

### Cleavage specificity and mechanism of 2’,5’-PEs

The 2’,5’-PEs studied here exclusively cleaved 2’,5’- and not 3’,5’-phosphodiester bonds. There was also a strong preference for cleavage of 2’,5’-oligoadenylates by NS2 and NS4b and, to a lesser extent, by VP3-CTD, whereas AKAP7 had similar activities against the different 2’,5’-linked pentamers of A, U, C and G. Therefore, although AKAP7 and VP3-CTD are not the mostly closely related 2’,5’-PEs, they can both cleave 2’,5’-oligoribonucleotides other than 2-5A (Fig. 2A and Table S2). Interestingly, OASs are 2’-nucleotidyl transferases that not only use ATP as substrates but can produce diverse molecules with 2’,5’ linkages. NAD^+^, tRNAs, A5’p_4_5’A, and mono- and poly-ADP-ribose are acceptors for addition of 2’,5’-AMP residues from ATP by OAS. Also, OAS can add other 2’-terminal ribo- and deoxy-nucleotide monophosphates to 2-5A (38–41). However, which, if any, of these alternative 2-5A-like molecules can be cleaved by 2’,5’-PEs remains to be determined. The cGAMP, a cyclic dinucleotide that activates STING, has one 2’,5’-linkage and one 3’,5’-linkage, but it is not cleaved by the 2’,5’-PEs examined here (Table S4). VP3 was phylogenetically distal and has the most distinct mechanism of 2-5A cleavage compared to all of the tested 2’,5’-PEs. It is also interesting to note that the two coronavirus 2’,5’-PEs (NS4b and NS2) are less closely related than NS2 is to the host enzyme, AKAP7 (Fig. 2A & Table S1). Our results suggest that the main, and perhaps only, function of these activities is to degrade 2-5A thus preventing RNase L activation and viral escape, or in the case of AKAP7 reducing cell and tissue damage from RNase L activity. These 2’,5’-PEs are also metal ion independent enzymes, as is RNase L (42).

The viral and mammalian 2’,5’-PEs produce cleavage products from trimer 2-5A (2’,5’- p_3_A_3_) with cyclic 2’,3’-phosphoryl groups, and not 2’,3’-OH termini. These conclusions are based on analysis of 2-5A cleavage products by two types of HPLC columns (Dionex and C18) and, importantly, by mass spectrometry. In contrast, our prior studies based on more limited analysis of the 2-5A cleavage products by one type of HPLC column (Dionex) misidentified these cleavage products of NS2, VP3-CTD, and AKAP7 as AMP and ATP (22, 25, 27).

Interestingly, mammalian and viral 2’,5’-PEs have activities highly similar to an invertebrate 2H-PE present in the oyster, Crassostrea gigas (43). The oyster enzyme has sequence similarity to AKAP7, is metal ion independent, cleaves 2’,5’- but not 3’,5’- linked oligonucleotides, and leaves cyclic 2’,3’-phosphate and 5’-OH termini on its products. It also degraded tri-phosphorylated 2-5A oligomers with multi-fold efficiency compared to the corresponding non-phosphorylated core 2-5A oligomers. Similarly, we observed RVA VP3-CTD degrades 5’-triphosphorylated-di-adenylate with 2’,3’ cyclic phosphoryl termini (p3ApA>p) preferentially compared to the non-phosphorylated di- adenylate (ApA) core molecule (Figs. S3 & S4). However, the function and role of 2-5A cleaving enzymes in invertebrates is still unknown.

It is also unknown if the 2’,3’-cyclic phosphates on 2-5A breakdown products generated by 2’,5’-PEs have cell signaling functions. However, small self RNAs with 2’,3’-cyclic phosphate termini (generated by RNase L) induced IFN-β expression through RIG-I, MDA5, and MAVS (44). Additionally, RNase L-cleaved RNA with 2’,3’-cyclic phosphates stimulated the NLRP3 inflammasome leading to IL-1β secretion (15). Also, during *Staphylococcus aureus* infections RNase T2 cleaves ssRNA producing purine-2’,3’- cyclic phosphate terminated oligonucleotides sensed by TLR8 (45). The mammalian enzyme USB1, another 2H-PE, also produces 2’,3’-phosphoryl temini during deadenylation of U6 snRNA, but it clearly differs from the 2’,5’-PEs because it cleaves 3’,5’-phosphodiester bonds (21).

### 2-5A catabolism during the IFN induced cellular responses to viral and host dsRNA

The ability of viruses to evade or antagonize the IFN response contributes to viral tropism and disease pathogenesis. Accordingly, viruses have evolved or acquired diverse strategies to overcome inhibition by type I and type III IFNs, both of which induce transcription of OAS genes (1, 2, 46). However, the precise cellular and molecular mechanisms by which viruses impede tissue specific host defenses leading to virus-induced pathology continue to be investigated. With regard to the OAS-RNase L pathway, it is the balance between 2-5A anabolic (OAS) and catabolic (e.g. 2’,5’-PEs and PDE12) (20) activities that determine whether virus replication is blocked by RNase L. For instance, RNase L fails to inhibit coronaviruses MHV and MERS-CoV, or rotaviruses, unless there is an inactivating mutation of their 2-5A degrading enzymes (NS2, NS4b, and VP3, respectively) (22, 24–26). In contrast, SARS-CoV-2, which lacks a gene for a similar protein, is inhibited by RNase L (4). In this context, enzymes that degrade 2-5A, such as PDE12, are drug targets in the hunt for broad spectrum antiviral agents (32, 47).

The viral enzymes NS2, NS4b and VP3-CTD are antagonists of innate immunity that support virus replication by eliminating 2-5A and preventing, or reducing, activation of RNase L by 2-5A (20, 22, 24–26). In contrast, mammalian AKAP7 is a nuclear 2’,5’-PE that does not affect viral replication, unless its nuclear localization signal peptide is deleted leading to cytoplasmic accumulation (27). A mutant AKAP7 deleted for its N-terminal nuclear localization signal peptide accumulates in the cytoplasm was able to rescue an NS2 mutant of MHV (22). While the function of the 2’,5’-oligonucleotide cleaving activity of AKAP7 is still unresolved, the phylogenetic tree suggests that the NS2 coronavirus proteins may have evolved from the AKAP7 catalytic domain (Fig. 2A).

Enzymes that degrade 2-5A have significance beyond antiviral innate immunity. Self-dsRNA activates the OAS-RNase L pathway leading in some circumstances to apoptosis (12, 13). In one example, mutation or inhibition of the dsRNA editing enzyme, ADAR1, leads to accumulation of self dsRNA activating OAS-RNase L leading to cell death, and PKR, inhibiting protein synthesis initiation (16, 48). In another instance, DNA methyltransferase inhibitors, e.g. 5-aza-cytidine, cause self-dsRNA accumulation from repetitive DNA elements leading to OAS-RNase L activation and apoptosis (17, 49). Thus, 2-5A is a secondary messenger for cytotoxic and antiviral activities of either non-self (viral) or self-dsRNA (host) whose levels must be tightly controlled to limit cytotoxicity while restricting viral spread. Our findings provide a mechanistic understanding of how 2’,5’-PEs regulate 2-5A levels among coronaviruses MHV, MERS-CoV, and group A rotaviruses and in mammalian cells through the activity of AKAP7 (22, 24, 25, 27), with implication for both control of virus replication and cellular responses to self-dsRNA. Furthermore, our study defines 2’,5’-PEs as a new sub-group within the 2H-PE superfamily that shares characteristic conserved sequence features of the superfamily but with specific and distinct biochemical cleavage activities. Knowledge derived from the study of these 2-5A degrading enzymes could lead to future avenues of antiviral drug development.

## MATERIAL AND METHODS

### cDNA cloning and plasmids

Human PDE12 cDNA was PCR amplified (using DNASU cDNA clone HsCD00296464 in vector pDONR221, GenBank: NM_177966.5) with forward primer 5’- TTCAA<u>gaattc</u>ATGTGGAGGCTCCCAGGCGC-3’ (with an EcoRI restriction site) and the reverse primer 5’- TTCAA<u>gtcgac</u>TCATTTCCATTTTAAATCACATACAAGTGCTATGTGATC-3’ (with a SalI restriction site). PDE12^E351A^ pGEX 6P-1 mutant (34) plasmid was constructed by MegaPrimer method (50) using mutagenic reverse primer 5’- GCGCGGTCAACCgCCTGCAAACAG-3’. The amplified wild type and mutant PDE12 cDNAs were cloned into plasmid pGEX-6P-1 (GE Healthcare, USA) at the EcoRI and SalI restriction site, sequenced and expressed in E. coli as glutathione *S*-transferase (GST) fusion proteins. To subclone the VP3 C-terminal domain (CTD) cDNA of rotavirus A strain RVA/Simian-tc/USA/RRV/1975/G3P (GenBank: EU636926.1) and its H718A mutant, we used codon-optimized constructs for expression in Sf9 insect cells (GenBank: KJ869109.1) (30) (gifts from Kristin Ogden, Vanderbilt University). The cDNAs were PCR amplified and cloned into plasmid pMAL-C5X at the XmnI (blunt cloned) and NcoI (sticky end) restriction site. Blunt end forward primer 5’- TACGCTGACGACCCCAACTACTTCATCG-3’ and reverse primer 5’- TTCAA<u>ccatgg</u>TTATTACTCGGACATGTCGAACACGGTGTCG-3’ with NcoI restriction site were used for VP3-CTD. The wild type and H718A RVA VP3-CTD proteins were expressed fused to maltose binding protein (MBP). Additional protein expression plasmid constructs were previously described with sequences originating from MERS-CoV (MBP-NS4b and its mutant MBP-NS4b^H182R^) (24); from MHV (MBP-MHV NS2 and its mutant MBP-NS2^H126R^) (22), and mouse AKAP7 and its mutant AKAP7^H93A; H185R^ (27).

### Protein expression and purification

Proteins were expressed from pGEX-6P-1 or pMAL constructs in E. coli strain BL21(DE3)/pLysS (Life Technologies, USA). Wild type and catalytically inactive mutants of AKAP7 and PDE12 were expressed as GST-fusion proteins and purification was performed by modification of a previous protocol (51). Single colonies were used to inoculate primary cultures which were subsequently used to seed secondary cultures grown to 0.6 OD (600 nm) in a shaking incubator at 37°C and 250 rpm. Cells were induced with 0.2 mM IPTG, for 16 h at 22°C. Induced cell pellets were re-suspended in buffer A [20 mM HEPES pH 7.5, 1 M KCl, 1 mM EDTA, 10% glycerol v/v, 5 mM DTT and EDTA-free Pierce™protease inhibitor (Thermo Scientific, USA)]. Pelleted cells were lysed by addition of 200 µg/ml lysozyme followed by sonication. Supernatants were collected after centrifugation at 12,000x g, 40 min, 4°C in a Beckman JA-17 rotor. Supernatants were added to Pierce™ Glutathione Agarose (Thermo scientific, USA) and incubated for 2 h at 4°C followed by washes with buffer A. Digestions to release the GST tag were performed with PreScission Protease (Cytiva, USA) in 50 mM Tris-HCL pH 7.5, 150 mM NaCl, 1 mM EDTA, and 1 mM DTT for 16 h at 4°C. Supernatants containing untagged protein were concentrated using Centriprep centrifugal filter devices (Millipore; molecular weight cutoff, 10 kDa) and loaded on superdex 75 column on an AKTA pure 25L protein purification system (GE Healthcare, USA) in 20 mM HEPES pH 7.5 150 mM NaCl, and 1 mM DTT. Wild type and mutant RVA VP3-CTD expressing bacterial culture growth and IPTG induction conditions were same as described above, except that growth media additionally included 2% glucose. Harvested bacterial cell pellets were suspended in Buffer B [20 mM Tris-HCl pH 7.4 with 200 mM NaCl, 1mM EDTA, 10 mM β-mercaptoethanol, EDTA free protease inhibitor (Pierce™ Protease inhibitor, Thermo Scientific, USA) and 10% glycerol] and lysed with lysozyme followed by sonication. Supernatants were incubated with Amylose resin (NEB, USA), washed three times with buffer followed by elution with 100 mM maltose. Proteins were concentrated using Centriprep centrifugal filter devices (Millipore; molecular weight cutoff, 10 kDa) and further purified using size exclusion chromatography (SEC) on an AKTA pure 25L protein purification system (GE Healthcare, USA) in buffer C (20 mM HEPES pH 7.5, 100 mM NaCl and 1 mM DTT). Wild type and catalytic mutants of NS4b, and MHV NS2 were purified as described previously (22, 24). In addition to inactive mutants, purified MBP protein was used as control in experiments with MBP fusion proteins. Protein concentrations were determined using Bio-Rad protein assay reagent (Bio-Rad, USA). All proteins were stored in Buffer C supplemented with 10% glycerol in -80°C.

### Synthesis and purification of 2-5A oligomers and other oligoribonucleotide substrates

2-5A or p_3_5’A(2’p5’A)_2_ (2’,5’-p_3_A_3_) was synthesized from ATP by using histidine-tagged porcine-OAS1 (52). The OAS was immobilized and activated with poly(I):poly(C)- agarose (53). Briefly, poly(I):poly(C)-agarose beads were washed with buffer D [10 mM HEPES pH 7.5, 1.5 mM Mg(CH_3_COO)_2_.4H_2_O, 50 mM KCl, 20% glycerol, and 7 mM β-mercaptoethanol]. Ten ml of beads were incubated with 10 mg of purified OAS protein for 2 h at 25°C with intermittent vortexing. Beads were washed three times with buffer D by centrifugation at 3000 g at 4°C for 30 min. Beads were suspended in the reaction mixtures containing 20 mM HEPES pH 7.5, 20 mM Mg(CH_3_COO)_2_.4H_2_O, 20 mM KCl, 1 mM EDTA and 10 mM ATP. The reaction mixtures were incubated in a shaking incubator set at 37°C, 120 rpm for 18 h. The supernatant was collected by centrifugation at 3000 g at 4°C for 30 min. The supernatant was heated at 95°C for 5 min and again centrifuged at 18,000 g for 15 min at 4°C to remove the precipitate. To isolate individual 2-5A oligomers, the supernatant containing crude, unfrationated 2-5A oligomers were run on an HPLC (1260 Infinity II Agilent technologies) equipped with a preparative Dionex column (BioLC^R^DNAPac^R^PA-100, 22 x 250 mm, Dionex, USA). Samples were injected and elution performed at a flow rate of 3 ml/min in a stepwise gradient of 10- 400 mM (0-120 min), 400-800 mM (121-125 min) and 10 mM (126-160 min) of NH_4_HCO_3_ buffer (pH 7.8). Fractions were collected, lyophilized and suspended in nuclease-free water.

RNA oligoribonucleotides (other than 2’,5’-p_3_A_3_) with 2’-5’ or 3’-5’ phosphodiester linkages were commercially purchased. Oligonucleotide substrates 5’- pA2’p5’A2’p5’A2’p5’A2’p5’A -3’, 5’-pU2’p5’U2’p5’U2’p5’U2’p5’U -3’, 5’-pG2’p5‘G2’p5‘G2’p5‘G2’p5‘G -3’, 5’-pA3’p5’A3’p5’A3’p5’A3’p5’A -3’, 5’-pU3’p5’U3’p5’U3’p5’U3’p5’U -3’ were purchased from Integrated DNA Technologies (IDT, USA) while 5’-pC2’p5’C2’p5’C2’p5’C2’p5’C -3’, 5’-pC3’p5’C3’p5’C3’p5’C3’p5’C -3’ and 5’-pA2’p5’A -3’ were purchased from ChemGenes Corporation (Wilmington, USA).

Penta-ribonucleotides substrates are shown as p5’(rN)_5_, where N represents A, U, G, or C nucleotide. A2’p5’A standard was prepared by incubating 5’pA2’p5’A with shrimp alkaline phosphatase (ThermoFisher, USA) as per manufacturer’s protocol. 2’,3’-cyclic-GMP-AMP (cGAMP), ATP, AMP and adenosine were obtained from Sigma Aldrich, USA.

### Phosphodiesterase activity assays

Ten µM of the substrates (with either 2’-5’ or 3’-5’ phosphodiester linkage) were incubated with 1 µM of enzyme. Final reactions were performed in 20 mM HEPES buffer pH 7.4, 1 mM DTT and 10 mM MgCl_2_ by incubating at 30°C for 1 h (or for the time indicated in the text). Where indicated, reactions were performed in the absence of MgCl_2_ with 2 mM EDTA added. Reactions were stopped by heating at 95°C for 5 min. Samples were centrifuged at 18,000 g for 15 min at 4 °C. Supernatants were collected and analyzed by HPLC. 2’,3’-cGAMP degradation assays were performed and analyzed using the same conditions as described above. In all experiments, substrates incubated under similar conditions in the absence of enzyme served as control.

### HPLC analysis and identification of products

The substrates and cleavage products were analyzed on a 1260 Infinity II Agilent technologies HPLC equipped with an Infinitylab Poroshell 120 C18 analytical column (Agilent technologies, 4.6 x 150 mm, 4µm). Eluent A was 50 mM ammonium phosphate buffer pH 6.8 and eluent B was 50% methanol in water. Five µl of processed samples were injected on the C18 column, at a flow rate of 1 ml/min and eluted with a linear gradient (0-40%) of eluent B over a period of 20 min, then 3 min 40% Eluent B, followed by equilibration to initial condition (100% Eluent A). The HPLC column was maintained at 40°C. Spectra were recorded at 256 nm. The products were identified either by comparing the elution time of known standards or by mass-spectrometry analysis. Alternatively, to test expanded substrate specificity, 10 µl of processed samples were injected on a Dionex DNAPac^R^PA-100 analytical column at a flow rate of 1 ml/min and eluted with a linear gradient of 10-800 mM of NH_4_HCO_3_ buffer (pH 7.8) over a period of 90 min, followed by 30 min equilibration to initial condition. Open Lab CDS software was used to analyze and calculate area under the peaks in HPLC spectra.

### Shrimp Alkaline phosphatase (SAP) mediated phosphorylation analysis

Purified substrates and cleavage product mixtures were dephosphorylated by incubating with SAP (ThermoFisher, USA) at 37°C for 1 h according to the manufacturer’s protocol. Samples were prepared for subsequent analysis as described above.

### Sample preparation for mass spectrometry

Desired peak fractions (including cleavage products of 2’,5’-p_3_A_3_) were collected by running samples on a Dionex DNAPac^R^PA-100 analytical column as described above. Collected peaks were subjected to acetone precipitation, supernatants containing cleavage products (from HPLC peak) were collected and lyophilized. Lyophilized samples were suspended in 1 mM NH_4_HCO_3_ buffer (pH 7.8) and used for mass spectrometry analysis.

### Mass spectrometry analysis of cleavage products

Prepared samples were subjected to mass spectrometry analysis. The LC/MS/MS analysis was carried out using a triple quadrupole tandem mass spectrometer (TSQ-Quantiva, Thermo Scientific, USA) equipped with an electrospray ionization (ESI) interface. The mass spectrometer was coupled to the outlet of HPLC system that consisted of an UHPLC system (Vanquish, Thermos Fisher Scientific, USA), including an auto sampler with refrigerated sample compartment and inline vacuum degasser. The Xcalibur software was used for data processing. The ESI mass spectrometric detection was performed in both the negative and positive ionizations, with ions spray voltage at 2.5kV, sheath gas at 35 Arb and Aux gas at 20 Arb. The ion transfer tube and vaporizer temperatures were set at 350°C and 250°C, respectively. The qualitative analysis was performed using full scan at the range from 200 to 1250 (m/z). Five µl extracted samples were injected on the C18 column (Gemini, 3 µm, 2 x 150 mm, Phenomenex, CA) with the flow rate of 0.3 ml/min at 45°C. Mobile phases were A (water containing 10 mM ammonium acetate and 20 mM ammonium hydroxide) and B (methanol containing 10 mM ammonium acetate and 20 mM ammonium hydroxide). Mobile phase B at 0% was used at 0-2 min, a linear gradient was used starting from 0% B to 100% B at 2-12 min, kept at 100% at 12-26 min, then from 100% B to 0% B at 26-27 min and kept at 0% B for 8 min. The peaks shown in full scans were processed to locate and identify the cleavage products of the 2’,5’-p_3_A_3_ substrate using the Xcalibur software v4.1. Standard adenosine, AMP, ATP and adenosine-2’,3’-cyclic monophosphate sodium salt were run for reference.

### Bioinformatic Analysis

The PDE domain sequences from different 2’,5’-PEs were used for creating a multiple sequence alignment using MAFFT version 7 (54) employing iterative refinement method E-INS-I (https://mafft.cbrc.jp/alignment/server/). The MAFFT sequence alignment result was downloaded in clustal format and visualized using Jalview 2.11.1.3 software. The sequence alignment was further processed on MAFFT server to calculate the phylogenetic tree using neighbor joining method and JTT substitution model and then visualized tree using Archaeopteryx.js software. The resultant fasta format output of MAFFT multiple sequence alignment was used to calculate the percentage of amino acid identity and similarity by Sequence Identity and Similarity (SIAS) tool with default parameters (http://imed.med.ucm.es/Tools/sias.html). The name, accession number and amino acid (aa) region of the aligned sequences are MHV NS2 (UniProtKB/Swiss-Prot: P19738.1, aa 41-135), Human coronavirus (HCoV) OC43 NS2 (GenBank: AAT84352.1, aa 43-138), Human enteric coronavirus (HECoV) NS2 (GenBank: ACJ35484.1, aa 39-140), Equine coronavirus (ECoV) NS2 (GenBank: ABP87988.1, aa 42-140), Middle East respiratory syndrome coronavirus (MERS-CoV) NS4b (GenBank: AFS88939.1, aa 87-191), Rat AKAP7 δ/γ (NCBI RefSeq: NP_001001801.1, aa 121-233), Mouse AKAP7 isoform-1 (NCBI RefSeq: NP_061217.3, aa 82-194), Human AKAP7 γ (NCBI RefSeq: NP_057461.2, aa 100-233), Human rotavirus group A (RVA) WA-VP3 (GenBank: AFR77808.1, aa 707-806), Simian rotavirus group A (RVA) SA11-N5 (GenBank: AFK09591.1, aa 707-808), Human rotavirus group B (RVB) Bang117 (GenBank: ADF57896.1, aa 655-750), Bat coronavirus (BtCoV) SC2013 NS4b (GenBank: AHY61340.1, aa 96-195) and Bat coronavirus (BtCoV) HKU5 NS4b (NCBI RefSeq: YP_001039965.1, aa 91-192).

## AVAILABILITY

Multiple sequence alignment software is available (https://mafft.cbrc.jp/alignment/server/). Alignment and phylogenetic tree construction tool is downloadable (https://www.jalview.org/). Sequence Identity and Similarity (SIAS) tool with default parameters is available (http://imed.med.ucm.es/Tools/sias.html).

## ACCESSION NUMBERS

The name and accession number of the aligned sequences are MHV NS2 (UniProtKB/Swiss-Prot: P19738.1), HCoV OC43 NS2 (GenBank: AAT84352.1), HECoV NS2 (GenBank: ACJ35484.1), ECoV NS2 (GenBank: ABP87988.1), MERS-CoV NS4b (GenBank: AFS88939.1), Rat AKAP7 δ/γ (NCBI RefSeq: NP_001001801.1), Mouse AKAP7 isoform-1 (NCBI RefSeq: NP_061217.3), Human AKAP7 γ (NCBI RefSeq: NP_057461.2), Human RVA WA VP3 (GenBank: AFR77808.1), Simian RVA SA11 N5 (GenBank: AFK09591.1), Human RVB Bang117 (GenBank: ADF57896.1), BtCoV SC2013 NS4b (GenBank: AHY61340.1) and BtCoV HKU5 NS4b (NCBI RefSeq: YP_001039965.1).

## ACKNOWLEDGEMENTS

We thank Dr. Renliang Zhang of Mass Spectrometry Core, Lerner Research Institute, Cleveland Clinic for performing LC/MS/MS and to Kristin Ogden (Vanderbilt University) for the rotavirus VP3-CTD cDNAs, and we thank Babal Kant Jha (Cleveland Clinic), Harpreet Kaur (Cleveland Clinic), Nikhil Bharambe (Case Western Reserve University) and Stephen A. Goldstein (University of Utah) for discussions.

## SUPPLEMENTARY FIGURES AND TABLES

**Figure S1. 2’,5’-p(A)_5_ degradation by MHV NS2 decreases in the presence of 2’,5’- p(U)_5_.** Substrate 2’,5’-p(A)_5_ was incubated with 0.2 mM of MHV-NS2 wild type protein in the absence or presence of indicated concentration of 2’,5’-p(U)_5_ at 30°C for 10 min. Samples were processed and analyzed by HPLC. 2’,5’-p(A)_5_ incubated under similar conditions in the absence of MHV-NS2 and 2’,5’-p(U)_5_ served as non-degraded control. Experiments were performed three times (n=3) and bars represent the standard error of mean. Statistical significance was calculated using unpaired t test (n=3; *, P value < 0.05; **, P < 0.005;***, P < 0.001; ns, not significant) in GraphPad Prism (9.0.0) software.

**Figure S2. MERS-NS4b degrades 2’,5’-p_3_A_3_ but not 2’,5’ linked ApA or pApA.** Substrate 2’,5’-p_3_A_3_ (A) was degraded in the presence of NS4b into p_3_A>p and ApA (B). Panels B, C, and D showed no appreciable decrease in amounts of ApA (area under the peak) after incubation for 1, 4 and 24 h, respectively. Panels E, F, G and H shows HPLC chromatograms of substrate pApA in the presence for NS4b at 0, 1, 4 and 24 h. (I) The table shows the amount of ApA or pApA degraded by NS4b as a function of time.

**Figure S3. Time-course of 2’,5’-p_3_A_3_ cleavage by RVA VP3-CTD.** Purified 2’,5’-p_3_A_3_ (200 µM) was incubated with RVA VP3-CTD (0.05 µM) at 30°C. Samples were collected at (A) 0 min, (B) 2 min, (C) 5 min, (D) 10 min, and (E) 30 min and analyzed by HPLC. The percent of substrate or products at indicated times were determined by calculating the area under the peaks on the HPLC chromatograms. Right hand side shows major and minor reactions proceeding at the indicated time points deduced from HPLC chromatogram analysis.

**Figure S4. Mechanism of 2’,5’-p_3_A_3_ cleavage by RVA VP3-CTD.** (A) The percentage of the substrate or the products at indicated times were determined by calculating the area under the peaks on the HPLC chromatograms obtained in experiment from figure S3. (B) Summary of major and minor reactions involved in cleavage of 2’,5’-p_3_A_3_ by VP3-CTD. The minor reaction cleavage of ApA to A>p and Ado is inferred from incubations performed at a 20-fold higher concentration of RVA VP3-CTD, 1 mM (Fig. 3D).

**Figure S5. RVA VP3-CTD degrades 2’,5’ linked di-adenylate.** (A) Substrate 2’,5’- pApA (A) was incubated with 1 mM of either (B) wild type RVA VP3-CTD or its mutant (C) RVA VP3 CTDH^718A^ at 30°C for 1 h. Samples were processed and analyzed by HPLC. Baseline buffer signal was subtracted from the samples.

**Figure S6. Time-course of 2’,5’-p_3_A_3_ cleavage by muAKAP7.** Purified 2’,5’-p_3_A_3_ (200 µM) was incubated with muAKAP7 (1 µM) at 30°C. Samples were collected at (A) 0 min, (B) 2 min, (C) 10 min, (D) 30 min and (E) 60 min and analyzed by HPLC. The peaks were identified by comparing the elution time of known standards. The percent of substrate or products at indicated times were determined by calculating the area under the peaks on the HPLC chromatograms. (F) Schematics showing cleavage of 2’,5’-p_3_A_3_ by muAKAP7. Reaction intermediate is shown in grey color.

**Table S1. Catalytic domain sequence identity and similarity analysis of viral and cellular 2’,5’-PEs.** Percent amino acid identity and similarity matrix is based on alignment in figure 2B. 2’,5’-PEs from mammals or mammalian viruses were used for alignment. Values are calculated using Sequence Identity and Similarity (SIAS) tool. Matrix values show percent identity (above diagonal) and similarity (below diagonal) between the corresponding pair of the proteins. Intragroup identity and similarity values are shaded in grey.

**Table S2. MERS-NS4b, MHV NS2, RVA VP3-CTD and muAKAP7 mediated degradation of 5’-phosphorylated, 2’-5’ or 3’-5’ linked penta-ribonucleotide substrates.** Ten µM of the indicated substrate was incubated with 1 µM of wild type or mutant 2’,5’-PE for 1 h at 30°C. Percent substrate degradation was calculated by measuring the area under the peaks in the HPLC chromatograms. Results were reproduced in at least two independent experiments.

**Table S3. MERS-NS4b mediated degradation of 5’-phosphorylated 2’-5’ or 3’-5’ linked penta-ribonucleotide substrates.** Ten µM of the indicated substrate was incubated with 3 µM of wild type or mutant MERS-NS4b for 1 h at 30°C. Percent substrate degradation was calculated by measuring the area under the peaks in the HPLC chromatograms. Results were reproduced in two independent experiments.

**Table S4. 2’,5’-PEs mediated degradation of 2’,3’-cGAMP and 2’,5’-p_3_A_3_.** Ten µM of the indicated substrate was incubated with 1 µM of wild type or mutant 2’,5’-PEs for 1 h at 30°C. Substrate without enzyme incubated under similar condition were used as un-degraded control. Percent substrate degradation was calculated by measuring the area under the peaks in the HPLC chromatograms. Results were reproduced in two independent experiments.

